# Characterization of HLA-restricted GAD65-specific CD8^+^ T cell responses in patients with GAD65 antibody-associated neurological disorders

**DOI:** 10.64898/2026.05.20.726502

**Authors:** Pei Shang, Benjamin DS Clarkson, Brittany L Overlee, Charles L Howe

## Abstract

**Background:** High-titer glutamic acid decarboxylase 65 (GAD65) antibodies are found in patients with GAD65 antibody-associated neurological disorders, including stiff-person syndrome (SPS), GAD65 cerebellar ataxia (CA), and GAD65 epilepsy. Given the intracellular localization of the antigenic target, a direct pathogenic role for GAD65 autoantibodies is unlikely. Instead, the autoantibody may be a biomarker for the existence of pathogenic anti-GAD65 autoreactive CD8^+^ T cells.

**Methods:** PBMC-derived dendritic cells (DCs) from 20 GAD65 antibody-positive patients with neurological manifestations (SPS, n=10; CA, n=7; epilepsy, n=3) and 15 healthy controls were pulsed with full-length GAD65 protein, full-length GAD67 protein, overlapping 15-mer peptide pools spanning GAD65 and GAD67, individual GAD65 15-mer peptides, or GAD65 9-mer peptides selected from predicted class I binding hotspots. T cell activation was quantified by flow cytometry-based activation-induced marker (AIM) assays using CD69 upregulation. Class I and class II HLA haplotypes were determined by high resolution typing. NetMHCpan v4.2c was used to generate residue-level peptide:HLA binding density maps across GAD65, and candidate 9-mers were validated for HLA binding by peptide:MHC monomer affinity testing. GAD65-peptide-HLA-restricted CD8^+^ T cells were identified by co-staining with two separately assembled tetramers carrying the same peptide:HLA complex on different fluorophores (APC and BV421), with double-positive events scored as antigen-specific. HLA-restricted cytotoxicity was measured by coculture of patient CD8^+^ T cells with GAD65-expressing HEK-293T cells reconstituted with defined HLA class I alleles using AAV-delivered Cre-dependent HLA-2A-eGFP cassettes.

**Results:** CD8^+^ T cells from GAD65 antibody-positive patients showed increased activation in response to DCs pulsed with full-length GAD65 relative to healthy controls (P=0.0157, Welch’s t-test), whereas responses to GAD67 did not differ significantly between groups. CD4^+^ T cells responded to both GAD65 (P=0.0004) and GAD67 (P=0.0051). Peptide pool screening of GAD65 identified discrete CD8^+^ and CD4^+^ immunogenic regions, with refinement using individual 15-mers localizing CD8^+^ activity to multiple subdomains within GAD65_(205-300)_, GAD65_(316-435)_, and GAD65_(447-520)_. HLA class I haplotyping in 16 non-Hispanic White GAD65^+^ patients revealed enrichment of HLA-B*08:01 (3.0-fold) and HLA-B*40:01 (4.1-fold) relative to USA NMDP European Caucasian reference frequencies (both BH q < 0.05), with suggestive enrichment of HLA-C*03:04 (2.9-fold; q = 0.09). Additionally, 44% of patients carried the HLA-A*01:01, HLA-B*08:01, and HLA-C*07:01 8.1 ancestral haplotype, approximately four-fold higher than the expected population frequency. Dual-fluorophore tetramer staining identified CD8^+^ T cells in GAD65^+^ subjects that bound a subset of HLA-A*11:01- and HLA-B*08:01-restricted GAD65 9-mers, with the clearest disease-skewed signals localized to GAD65_(213-221)_, GAD65_(257-265)_, and GAD65_(529-537)_. In an HLA-reconstituted target-cell killing assay, CD8^+^ T cells from an HLA-B*08:01-positive GAD65^+^ patient mediated antigen- and HLA-restricted depletion of GAD65-expressing HEK-293T cells, with HLA-restricted target loss also observed in single-donor experiments across additional HLA-A, -B, and -C contexts.

**Conclusions:** Patients with GAD65 antibody-associated neurological disorders harbor circulating CD8^+^ T cells that recognize discrete HLA class I-restricted GAD65 peptides and that are capable of cytotoxicity against GAD65-expressing HLA-matched target cells. We characterize the immunogenetic and cellular features of class I-restricted CD8^+^ T cell responses in GAD65 autoimmunity, including overrepresentation of the 8.1 ancestral haplotype, complementing the historical focus on antibodies and CD4^+^ T cell help, and we provide a panel of validated GAD65 peptide:HLA tetramers for prospective isolation, clonotypic analysis, and longitudinal monitoring of candidate pathogenic CD8^+^ T cell populations across the GAD65 antibody-associated neurological disease spectrum.

## Introduction

Glutamic acid decarboxylase (GAD) is the rate-limiting enzyme in synthesis of the inhibitory neurotransmitter gamma-aminobutyric acid (GABA) through decarboxylation of L-glutamate ^1^. The two isoforms of GAD are GAD65 (encoded by GAD2 on human chromosome 10) and GAD67 (encoded by GAD1 on human chromosome 2), named according to molecular weight ^2^. The two isoforms share 75% homology, predominantly in the middle segment and C terminus ^3^. GAD65 is enriched in axon terminals due to N-terminal domain anchoring to the membrane of presynaptic vesicles, whereas GAD67 is mainly located in the neuronal cytoplasm ^4^. GAD67 is constitutively active and responsible for over 90% of GABA production in the central nervous system (CNS) ^5^, while GAD65 is responsible for activation and augmentation of GABA levels for rapid modulation of inhibitory neurotransmission ^6^. Both isoforms are expressed in GABAergic inhibitory neurons as well as in pancreatic islet cells, and antibodies against GAD65 have been identified in both patients with type I diabetes (T1D) and patients with neurological disorders ^7^.

While GAD antibodies are detected at low levels in healthy individuals and patients with numerous neurological disorders, high-titer GAD antibodies are associated with a defined spectrum of neuronal excitability syndromes, including stiff-person syndrome (SPS) ^8^, cerebellar ataxia (CA), epilepsy, and limbic encephalitis ^9^. A retrospective review of 212 GAD65 patients with neurological manifestations from 2003 to 2018 showed that 50% of patients had SPS spectrum disorders, 43% had cerebellar ataxia, 29% had autoimmune epilepsy, and 16% had limbic encephalitis ^10^. SPS predominantly presents in adult women and is characterized by stiffness, muscle spasms, hyperexcitability, and abnormal gait with falls. GAD antibody-positive cerebellar ataxia often co-presents with SPS or T1D and is marked by gait and limb ataxia, dysarthria, nystagmus, and ocular motor dysfunction. GAD65-associated limbic encephalitis presents with psychiatric symptoms, memory deficits, and seizures, similar to other autoimmune encephalopathy patterns ^11^. Most GAD65 antibody-positive patients with neurological manifestations are pharmaco-resistant and are treated palliatively.

GAD65 immunoglobulins in both T1D and GAD antibody-positive neurological diseases are predominantly the IgG1 subtype ^12, 13^. GAD antibodies isolated from patients with SPS recognize linear epitopes in N-terminal, C-terminal, and protein-lipid binding domains ^3, 14^. In contrast, GAD antibodies in patients with T1D primarily recognize conformational epitopes at the C-terminal and protein-lipid binding domains ^15^, suggesting immunodominant epitope differences between these diseases. Patients with limbic encephalitis may exhibit more antibody binding to the N-terminal domain of GAD65 compared to antibodies from patients with SPS, CA, and epilepsy, in which antibodies exhibit more binding to C-terminal domains ^16^. However, other findings suggest that these differences are not specific ^17^. In clinical practice, anti-GAD65 antibodies have greater diagnostic specificity and sensitivity when compared to anti-GAD67 antibodies ^18^. The presence of antibodies to GAD65 in patients with SPS and other neurological manifestations has led to the hypothesis that the antibodies directly mediate loss of inhibitory neurotransmission in the cortex, brainstem or spinal cord and induce stiffness, rigidity, or startle responses in patients, despite the intracellular location of the antigen. These observations have motivated numerous clinical trials targeting the antibody and antibody-producing cells in patients with SPS and other neurological manifestations of anti-GAD autoimmunity ^19^. However, a direct pathogenic role for anti-GAD antibodies in these patients remains controversial and unproven ^20^.

Outside of a direct pathogenic mechanism, the presence of high avidity anti-GAD65 autoantibodies in patients with neurological manifestations ^21^ indicates a role for anti-GAD CD4^+^ T cells, based on T follicular helper cell support of B cell somatic hypermutation and antigen-driven immunoglobulin affinity maturation ^22, 23^. HLA class II-restricted clonally expanded GAD65 peptide-specific CD4^+^ T cells have been detected in CSF and blood in patients with SPS ^24^ and 80% of SPS patients with autoimmune comorbidities such as thyroiditis, T1D, pernicious anemia, and vitiligo show strong associations with DR and DQ alleles ^14, 25^. It is notable that transgenic mice expressing GAD65-specific TCRs on CD4^+^ T cells exhibited a spontaneous lethal encephalomyelitis syndrome ^26^ and adoptive transfer of GAD65-specific CD4^+^ T cells into non-diabetic NOD/SCID mice induced insulitis ^27^. However, despite this evidence, it is unclear whether anti-GAD65 CD4^+^ T cells directly contribute to anti-GAD autoimmunity in humans with SPS and other neurological manifestations associated with GAD65 antibodies.

Several observations support a role for cytotoxic CD8^+^ T cells in anti-GAD65 autoimmunity. Autopsy and surgical neuropathology studies in SPS and GAD-associated epilepsy demonstrate infiltrating CD8^+^ cytotoxic T cells within clinically relevant CNS lesions, consistent with cell-mediated injury ^28, 29^. In parallel, in human T1D, antigen-specific CD8^+^ T cells targeting GAD65 have been directly detected using HLA class I tetramers ^30^, and HLA-A*02:01-restricted CD8^+^ T-cell clones recognizing the immunogenic region around GAD65_(114-123)_ can kill human islet cells, supporting natural processing/presentation of GAD65 class I epitopes in human tissue ^31^. Nonetheless, compared with the T1D-focused literature, the specificity, breadth, and antigen-processing requirements of GAD65-directed CD8^+^ T cell responses in SPS and other GAD65 antibody-associated neurological disorders remain incompletely defined, motivating systematic evaluation of CD8^+^ responses to GAD65 in these conditions. To address this gap, we used an activation-induced marker (AIM) assay to measure CD4^+^ and CD8^+^ T cell responses to both full-length GAD65 antigen and to a library of overlapping 15-mer peptides derived from GAD65. We also identified candidate immunodominant 9-mer peptides and measured the frequency of GAD65 peptide:MHC class I specific CD8^+^ T cells among PBMCs collected from patients with GAD65 antibody-associated neurological disorders. Finally, we tested the capacity of GAD65-specific CD8^+^ T cells to injure GAD65-expressing HLA class I-matched target cells.

## Material and Methods

### Patient cohort

All patients consented to participate in the study (IRB#23-003442). The cohort comprised 20 patients with GAD65 antibody-associated neurological disorders (SPS, n=10; cerebellar ataxia, n=7; epilepsy, n=3) and 15 healthy controls. Serum GAD65 antibody titers in the patient cohort ranged from 25 to 1707 nmol/L, all above the clinical cutoff of 20 nmol/L. Patients received variable immunotherapy in the 6 months preceding PBMC collection, including intravenous methylprednisolone, intravenous immunoglobulin, plasma exchange, rituximab, mycophenolate mofetil, methotrexate, azathioprine, or no treatment (Table 1). Treatments were not impacted or adjusted as a result of this study. Per-figure subject inclusion varied with specimen availability, recruitment timing, and HLA matching.

**Table 1.**
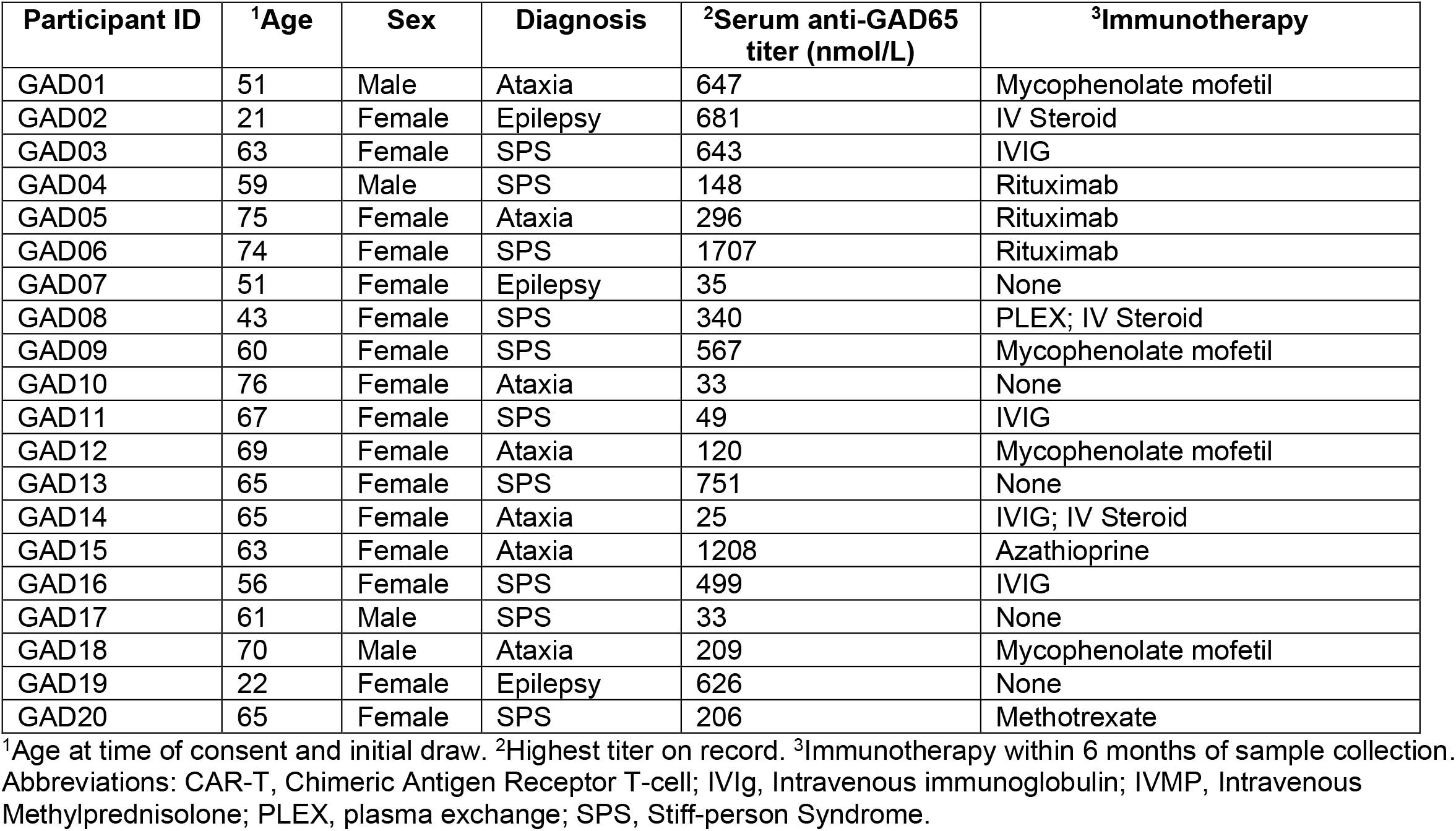
GAD65^+^ patient demographics.

### Isolation and cryopreservation of peripheral blood mononuclear cells from patient blood

Blood from patients and healthy donors was collected in EDTA tubes and PBMCs were isolated using Leucosep centrifuge tubes (VWR) as described in our previous publication ^32^. The bottom chamber of the Leucosep tube was filled with Lymphoprep (Stemcell Tech) and top chamber overlaid with fresh patient whole blood. Tubes were spun at 800g for 15 min at room temp with no brake. Buffy coat was collected and washed once with RPMI 1640 media. Cell pellet was resuspended and counted using a flow cytometer (Attune NxT, Thermofisher). PBMCs were frozen down at 5×10^6 cells/vial in freeze media consisting of 10% DMSO, 45% Human serum (Sigma) in RPMI (Thermofisher).

### Dendritic cell differentiation

Cryopreserved PBMCs were washed in 15% FBS RPMI 1640 media, spun at 400g for 5 mins and plated at 2 × 10^6^ cells per well of 6 well dishes in RPMI 1640 containing GM-CSF (50 ng/mL; *Peprotech, Catalog:300-03*) and IL-4 (20 ng/ml; *Peprotech, Catalog:200-04*) for dendritic cell (DC) differentiation. Remaining cells were rested with IL-7 (10 ng/ml; *Peprotech, Catalog:200-07*) to maintain the T cell population.

### Antigen pulsing

DCs and T cells were cocultured with autoantigens including full-length GAD65 (FL-GAD65; *Origene, Catalog: TP325984*) and GAD67 (FL-GAD67), GAD peptides (226 overlapping peptides o*rdered from GeneScript*), or peptide pools from CMV, EBV, and Influenza virus CEF (*AnaSpec, Catalog:AS-61036-003*) as positive control. GAD peptides were 15 amino acids long with 10 amino acids overlapping adjacent peptides. All antigens were reconstituted at 10 mg/ml. For 15mer peptide pools, we combined up to 10 peptides at a final concentration of 1 mg/ml. Peptide pools were used as antigens 10 µg/well (100 µg/mL) to stimulate the cocultured DCs and T cells on 96-well plate. In some AIM assays, single or pooled 9mer peptides were similarly used at 100 µg/mL final concentration. Co-cultured DCs and TCs were treated with antigens for 3 days prior to analysis by flow cytometry with CD3, CD4, CD8, and CD69.

### Flow cytometry

Antibodies against human lymphocyte markers CD3 (PerCP-Cy5.5) (*Biolegend, Catalog:300328*), CD8 (PE) (*Biolegend, Catalog:300908*), CD4 (APC) (*Biolegend, Catalog:317416*), and the T cell activation marker CD69 (FITC) (*Biolegend, Catalog:310904*) were used as antigen-induced marker panel in our study. Prior to staining, cells were washed twice in buffer comprised of calcium magnesium-free PBS with 2 mM EDTA, 5% FBS, 1% BSA and 0.01% sodium azide. Cells were then labeled with antibodies for 30 min on ice with agitation. Cells were washed 3 times with buffer and acquired on an Attune NxT Flow cytometer (ThermoFisher). FCS files were exported and analyzed in FlowJo software version 10.4.2 (Tree Star Inc.). Gating thresholds for positive/negative events were set using fluorescence-minus-one (FMO) and isotype control samples.

### Molecular HLA Haplotyping

HLA typing was performed using PBMCs from selected GAD65 and healthy donors. Genomic DNA was extracted from PBMCs by conventional techniques. Both class I and class II HLA typing was performed with high resolution (4-digit resolution), HLA-A, -B, -C, -DRB1, and -DQB1 (Creative Biosciences, CD Genomics).

### HLA class I peptide hotspot analysis

We predicted peptide:HLA binding across the human GAD65 protein using NetMHCpan v4.2c and extracted the peptide start position, peptide sequence, and eluted ligand rank score (EL_rank) for each predicted binder. HLA class I allotypes detected in GAD65^+^ subjects were used for the analysis. To generate a residue-level binding hotspot map, each peptide was assigned a binding weight based on its EL_rank, using a monotonic transformation in which lower ranks contributed larger weights [max (0, −log _10_(EL_rank/2))]. This weight was distributed evenly across all amino acid positions spanned by the peptide by dividing the peptide weight by peptide length, and the per-residue contributions from all overlapping peptides were summed to generate a combined hotspot score at each residue. The resulting residue-wise score reflects the local density and breadth of predicted HLA presentation along the GAD65 sequence, with higher values indicating regions enriched for overlapping peptides with stronger predicted binding across the analyzed HLA allotypes.

### Peptide MHC Monomer Affinity Tests

Peptide MHC Monomers were generated with peptides of interests (peptides inducing T cell activation and predicted to have high affinity to patients’ HLA alleles) or positive control peptides using HLA easYmers: *HLA*A02:01(Immundex, Cat:U-LWBM), HLA*A03:01(Immundex, Cat:1016-01), HLA*A11:01(Immundex, Cat:U-LWDM), HLA*B07:02(Immundex, Cat:1048-01), HLA*B08:01(Immundex, Cat:1050-01), HLA*B18:01(Immundex, Cat:1066-01), HLA*B40:01(Immundex, Cat:U-LVKM), HLA*C 03:03(Immundex, Cat:1116-01), HLA*C 03:04(Immundex, Cat:U-LWSM), HLA*C 07:01 (Immundex, Cat:1121-01), HLA*C 07:02(Immundex, Cat:U-LVYM)*. For each test, 3 µL of GAD65 peptides (75 µM) were mixed with 12 µL easYmer, 12 µL loading buffer and 45 µL ddH_2_O and incubated for 48 hours at 18 °C to generate 72 µL 500 nM monomers. Each monomer solution (6 µL) was then added into 69 µL dilution buffer (5% glycerol PBS) to generate 40 nM monomers for affinity tests. For affinity tests, 40 nM monomers were serially diluted to make 13.3 nM, 4.4 nM, and 1.5 nM solutions. Streptavidin beads (6-8 µm, Spherotech, Cat:SVP-60-5, 1:45 dilution) were incubated with monomers at 37°C for 60 mins, washed, centrifuged (700xg for 3 min), and incubated with β2m monoclonal antibody BBM.1-PE (Santa Cruz Cat# sc-13565 PE, 1:200 dilution) for 30 mins. Beads were then washed 3 times and acquired an Attune NxT Flow Cytometer to measure mean fluorescence intensity. All peptide affinity to HLA binding parameters were compared with positive and negative controls and subclassified into strong, intermediate, low, and no-binders.

### Tetramer Staining

For monomers validated to be strong or intermediate binders, 25 ul monomers (500nM) were incubated with 20ul Streptavidin-APC *(BD; Cat#554067; 0*.*2mg/ml)* or 40ul Streptavidin-BV421 *(BD; Cat# 563259; 0*.*1mg/ml)* separately at 4°C for 60 mins to make tetramers. Empty MHC monomers were cultured with Streptavidin-FITC *(BD; Cat#554060; 0*.*5mg/ml)* at 4°C for 60 mins as negative controls. Cryopreserved PBMCs from patients with GAD65 AbAD and HCs were thawed and rested for 1 hour to recover surface expression of TCR epitopes. PBMCs were incubated with 30 nM APC-labeled—tetramers and BV-421-labeled—tetramers complexed to GAD65 peptides, along with FITC-labled-empty (negative control) tetramers. The incubation was performed in fluorescence-activated cell sorting (FACS) buffer (PBS containing 3% bovine serum) at room temperature for 20 min in dark, followed by washing and further surface staining with PE-labeled mAb to CD8 (eBioscience), PerCP-Cy5.5-labeled mAb to CD3 (eBioscience) for 30 min at 4°C. FITC-negative, dual tetramer^+^ cells were analyzed within a CD8^+^ cell gate. Cells were acquired with Attune NxT Flow Cytometer (ThermoFisher) and analyzed in FlowJo software version 10.4.2 (Tree Star Inc.).

### Coculture Assays

HEK293T cells were transfected with LV.TagBFP2.EF1A.hGAD2-T2A-iCre or LV[Exp]TagBFP2.EF1A.iCre and purified by flow sorting BFP^+^ cells. These cells were then expanded and purity confirmed by flow cytometric analysis before cryopreservation. Cryo-preserved cells were thawed and seeded in a T-175 flask with HEK cell media (500 mL DMEM, 3%FBS, 1% Pen/Strep) for 3 days. Cells were then harvested and pretreated with 1 µM dorsomorphin for 24 hours before transfection with AAV[FLEX]-HLA-eGFP vector (e.g., AAV-CAG-DIO-HLAB08:01-P2A-eGFP-WPRE) or AAV-CAG-DIO-eGFP-WPRE empty vector control (at 2x10^9^ gc/mL) in a 96-well glass bottom plate (CellVis). HEK293T target cells were monitored for 1-3 days to confirm eGFP expression, after which HLA-matched patient and control PBMCs were thawed and rested in 10 ng/ml IL-2/IL-7 24 hours. CD8^+^ T cells were purified by magnetic activated cell sorting using the EasySep Human CD8^+^ T cell isolation kit (StemCell Technologies) before coculture with HEK293T target cells. Automated live cell imaging was performed using an SX3 IncuCyte (Sartorius). Wells were imaged every 4 hours from t = −24 h through endpoint. eGFP^+^ area and phase-contrast confluence were quantified in IncuCyte software, and downstream analysis is described in the Statistics section. At selected endpoints, cocultures were harvested for flow cytometric analysis of HEK target survival (percent eGFP^+^ in CD3-cells, normalized to no-T-cell control) and T cell activation (CD69 on CD3^+^CD8^+^).

### Statistics and Data Analysis

#### AIM activation score calculation

For each subject and condition, the percentage of CD69^+^ cells within the CD8^+^ or CD4^+^ T cell compartment was measured by flow cytometry and background-corrected by subtracting the unstimulated condition for the same subject. Because background-corrected frequencies can be negative and span multiple orders of magnitude, values were transformed using the inverse hyperbolic sine, asinh(x), prior to scaling ^33^. Scaled activation scores were computed using the median absolute deviation (MAD), defined as MAD(x) = median(|x − median(x)|) ^34^, according to:

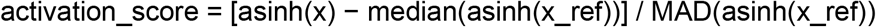

where x_ref is the reference distribution chosen for the analysis context. For peptide-level responder calling, x_ref was the control distribution for the same peptide; for subject-centered analyses, x_ref was the unstimulated condition for the same subject. A positive activation score indicates an increase in CD69^+^ frequency relative to baseline; a value of 0 indicates no change.

#### Responder thresholding

For peptide- and tetramer-level responder calling, a control-based positivity threshold was defined separately for each peptide or peptide:HLA complex j as T_j = median(CON_j) ^+^ 1.5 × MAD(CON_j), where CON_j is the control distribution for that peptide or tetramer. Subjects were classified as responders for j if their activation score exceeded T_j. Responder proportions were computed within each group for each peptide or tetramer.

#### Group comparisons

Continuous activation scores were compared between GAD65^+^ subjects and controls using unpaired t-tests with Welch’s correction for unequal variance applied to asinh-transformed scores. Where the question of interest was whether scores within a defined subset differed from baseline rather than between groups, one-sample Wilcoxon signed-rank tests were applied to subject-centered scores against a null median of 0. Tetramer binding scores were compared between groups using Wilcoxon rank-sum tests within HLA-matched subsets where both groups had n≥3.

#### HLA enrichment

HLA class I allele frequencies in GAD65^+^ patients were compared to reference frequencies from the USA NMDP European Caucasian panel (n = 1,242,890 typed individuals) in the Allele Frequency Net Database (AFND, allelefrequencies.net) ^35^. To ensure ancestral compatibility with the reference panel, HLA enrichment analyses were restricted to the 16 GAD65^+^ patients self-identifying as non-Hispanic White; two GAD65^+^ patients of Hispanic/Latino background were excluded from the population reference comparison. A non-GAD65^+^ control group of 12 individuals who self-identified as non-Hispanic White was used for comparison. For each allele observed in the patient cohort, observed counts were tested against reference frequencies using two-sided exact binomial tests and Pearson chi-square statistics. P-values were adjusted across all observed alleles using the Benjamini-Hochberg false discovery rate procedure (BH q-values). Per-locus deviation of the patient HLA distribution from reference was assessed by Pearson chi-square goodness-of-fit testing, with reference alleles not observed in patients pooled into an ‘other’ category. Alleles observed in a single patient chromosome were noted descriptively but were not included in formal significance testing because per-allele frequency estimates from n=1 are unstable. Joint carriership of the HLA-A*01:01, HLA-B*08:01, HLA-C*07:01 8.1 ancestral haplotype was scored when a patient carried at least one copy of each constituent allele and was tested against the published carrier frequency for this haplotype in US European populations (approximately 12%, corresponding to a haplotype frequency of approximately 6%) ^36^ using a two-sided exact binomial test.

#### Target-cell killing analysis

For HLA-reconstituted HEK-293T killing assays, endpoint flow cytometric quantification of the percentage of eGFP^+^ cells within the CD3-HEK gate was normalized to the no-T-cell condition. Differences across HLA condition and effector source (patient vs. control) were assessed by two-way ANOVA with Sidak’s correction for pairwise comparisons. Live-cell imaging data were acquired at fixed intervals on an IncuCyte SX3 (Sartorius); per-well eGFP^+^ area was normalized to phase-contrast confluence to control for variation in target-cell number, and the resulting GFP^+^ area per confluence was further normalized to the baseline value at coculture initiation. Each condition was tested with a minimum of 6 replicate wells; per-condition replicate counts are given in the figure legends.

#### Multiple testing

Where multiple parallel peptide-, tetramer-, or subgroup-level tests were performed, raw p-values were adjusted using the Benjamini-Hochberg false discovery rate procedure, and adjusted q-values are reported alongside nominal p-values.

#### General reporting

All analyses followed the guidelines of Curran-Everett ^37^. Normality assumptions were assessed by Shapiro-Wilk testing where applicable; for activation scores, non-parametric tests were used by default given the bounded, transformed nature of the scaled scores. P-values are reported as two-sided. Figures show individual-subject values with summary bars (mean ± 95% CI as indicated); dashed horizontal lines on AIM plots indicate an activation score of 0; black horizontal bars on peptide-level plots indicate the responder threshold (control median ^+^ 1.5 × MAD).

## Results

### Full-length GAD65 and GAD67 proteins elicit antigen-specific T cell activation in GAD65^+^ patients

We investigated antigen-specific CD8^+^ and CD4^+^ T cell responses using activation-induced marker (AIM) assays at 72 hr after co-culture with autologous PBMC-derived dendritic cells (DCs) exposed to full-length GAD65 or GAD67 proteins. Surface levels of CD69 were assessed by flow cytometry and the frequency of CD8^+^CD69^+^ or CD4^+^CD69^+^ cells for each subject was normalized to the basal CD69^+^ frequency in co-cultures without antigen exposure. An activation score was calculated using an inverse hyperbolic sine (asinh) transformation ^33^ scaled to the median absolute deviation ^34^ for each condition to accommodate negative values associated with the background correction. A positive activation score indicates an overall increase in the frequency of CD69^+^ T cells relative to unstimulated. The GAD65-positive patient cohort (n=10) showed a significant increase in CD8^+^ T cell activation in response to DCs fed whole GAD65 antigen relative to the control cohort (n=9) (P=0.0157, t-test with Welch’s correction) (Figure 1A). In contrast, both GAD65^+^ subjects and controls showed similar activation in response to DCs fed whole GAD67 protein (P=0.1299) (Figure 1B). CD4^+^ T cells from GAD65^+^ subjects responded more robustly than controls to both GAD65 (P=0.0004) (Figure 1C) and GAD67 (P=0.0051) (Figure 1D), consistent with previous reports. We verified the overall T cell responsiveness in the subjects by measuring activation induced by DCs loaded with a cocktail of immunogenic peptides from human cytomegalovirus, Epstein-Barr virus, and influenza virus (CEF). As expected for this cocktail, CD8^+^ T cell responses (Figure 1E) were larger than CD4^+^ T cell responses (Figure 1F), but both controls and GAD65^+^ subjects showed equivalent activation profiles.

**Figure 1.**
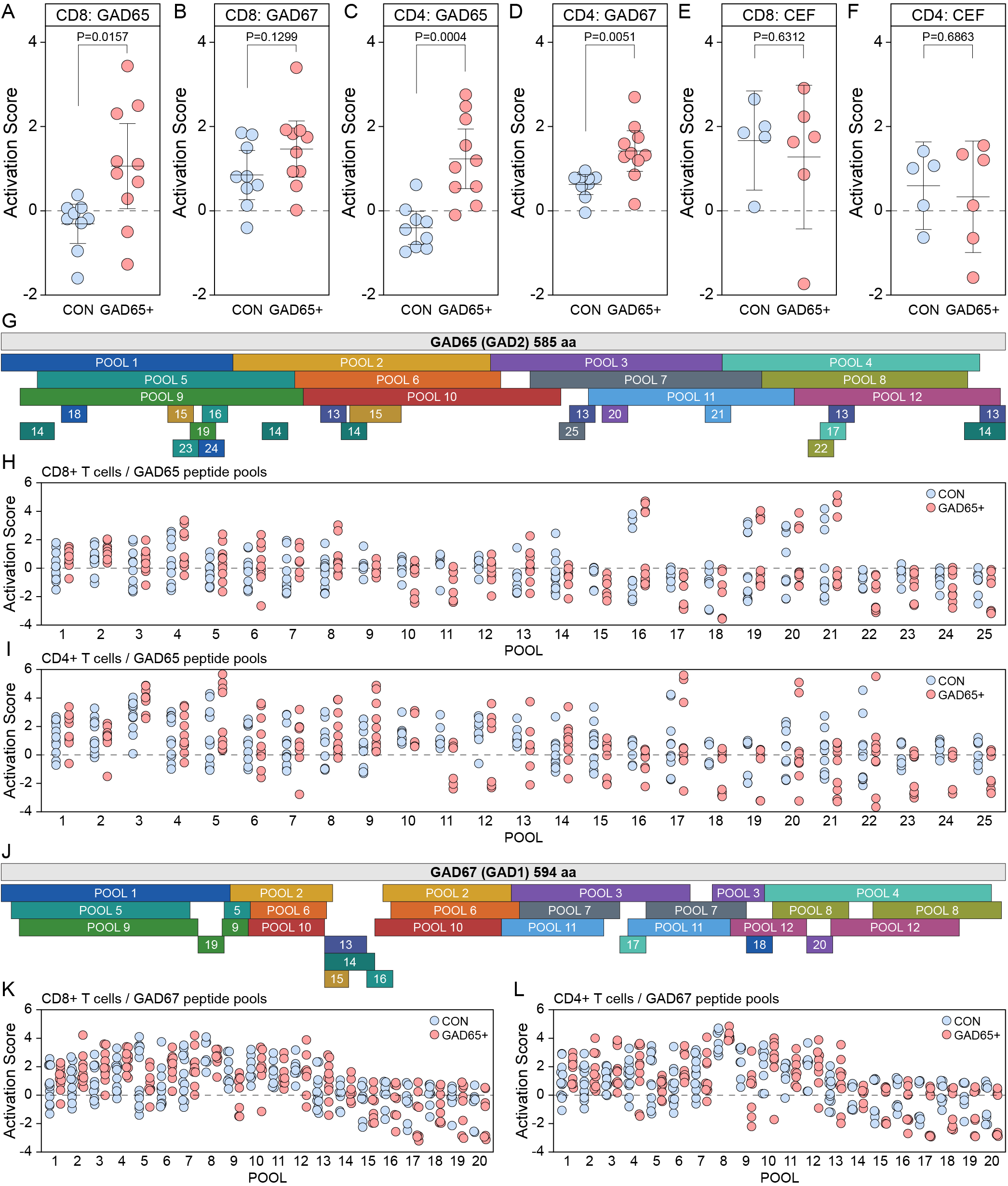
Full-length GAD proteins and overlapping peptide pools reveal antigen-specific T cell activation in GAD65^+^ subjects. **A-F**) Activation-induced marker (AIM) responses of CD8^+^ and CD4^+^ T cells after 72 h co-culture with autologous PBMC-derived dendritic cells loaded with full-length GAD65, full-length GAD67, or a control CEF peptide cocktail. Responses are shown as activation scores derived from background-corrected CD69 frequencies using asinh transformation and MAD scaling. Each point represents one subject; error bars indicate mean ± 95%CI; P-values show Welch’s t-test between GAD65^+^ and CON. Dashed horizontal lines indicate an activation score of 0. Subjects used for the experiments in panels A–F (n=10 GAD65^+^, n=9 controls) were those enrolled and processed at the time of the AIM experiment; per-figure inclusion across the study varied as described in methods. **G, J**) Schematic maps of overlapping 15-mer peptide pools spanning the full-length GAD65 (585 aa; **G**) and GAD67 (594 aa; **J**) proteins. Adjacent peptides overlap by 10 aa. For screening, peptides were grouped into larger primary pools and smaller secondary pools, with additional secondary pools created to accommodate peptides with lower aqueous solubility. Numbered boxes indicate the secondary pools used for deconvolution. **H**) CD8^+^ and **I**) CD4^+^ AIM activation scores in response to GAD65 peptide pools. Each point represents one subject tested against the indicated pool. Several GAD65 regions elicited activation in subsets of subjects, including responses detected in secondary pools that were not fully captured by the corresponding larger parent pools, consistent with localized immunodominant domains and cell type-specific recognition patterns. **K**) CD8^+^ and **L**) CD4^+^ AIM activation scores in response to GAD67 peptide pools. As for GAD65, peptide pool screening identified discrete domains of GAD67 capable of activating both CD8^+^ and CD4^+^ T cells. Blue circles, controls; red circles, GAD65^+^ subjects.

### Identification of immunodominant domains in GAD65 and GAD67 proteins

To map the specific antigenic regions within the full-length GAD65 and GAD67 proteins, we divided each protein into overlapping 15-mer peptides covering the entire 585 amino acid sequence of GAD65 and the 594 amino acids of GAD67. Each peptide overlapped with the adjacent sequence by 10 amino acids to ensure comprehensive coverage. For practical screening, we grouped these peptides into pools of up to 10 peptides each. Peptides that showed low solubility in double-distilled water and peptides with lower overall aqueous solubility were grouped and tested separately, resulting in 12 large pools and 10 secondary pools for GAD65 (Figure 1G) and 12 large pools and 8 secondary pools for GAD67 (Figure 1J). Using these peptide pools in AIM assays we observed a pattern of GAD65 domains that activated CD8^+^ T cells in both controls and GAD65^+^ subjects (Figure 1H). For example, small peptide pools 16 and 19 induced activation in a subset of subjects, as did larger pool 5, while the surrounding regions of pools 1 and 9 did not. Likewise, small pools 20 and 21 induced activation that was not captured in pool 3, 7, or 11. Finally, pools 4 and 8 but not pool 12 induced activation and the distal terminus of GAD65 present in pool 13 exhibited some activation that was not present in pool 14. Different pools conferred activation in CD4^+^ T cells relative to CD8^+^ cells (Figure 1I); for example, pool 9 induced strong activation in many subjects but pools 16 and 19 did not. Using the same strategy, we also identified domains in GAD67 that induced CD8^+^ (Figure 1K) and CD4^+^ (Figure 1L) activation.

On the basis of these findings, we tested individual 15-mers selected from candidate immunogenic regions (Figure 2). This assay identified multiple domains from GAD65_(205-300)_, GAD65_(316-435)_, and GAD65_(447-520)_ that activated CD8^+^ T cells in both controls and GAD65^+^ subjects (Figure 2A), as well as subdomains such as GAD65_(241-255)_ and GAD65_(251-265)_ that were skewed toward GAD65^+^ subjects. Of note, domains such as GAD65_(451-465)_ induced larger responses in controls relative to GAD65^+^ subjects. Similar mapping revealed CD4^+^ T cell hotspots in GAD65 (Figure 2B). Further refinement with pools of 9-20-mers covering GAD65_(211-225)_, GAD65_(240-285)_, GAD65_(446-470)_, and GAD65_(501-516)_ isolated two subdomains that strongly activated both healthy controls and GAD65^+^ subjects (Figure 3, POOL2 and POOL6) and one subdomain at GAD65_(501-516)_ that skewed toward more activation in GAD65^+^ subjects (Figure 3, POOL8). The variable responses observed with different peptide lengths and coverage prompted us to examine the distribution of HLA class I haplotypes across the GAD65^+^ cohort (Table 2). HLA class I haplotyping was completed in 18 of 20 GAD65^+^ patients. To ensure ancestral compatibility with the USA NMDP European Caucasian reference panel, enrichment analyses were restricted to the 16 non-Hispanic White patients (32 chromosomes per locus); two Hispanic/Latino patients were excluded from this comparison. Two alleles were enriched in GAD65^+^ patients at BH q < 0.05: HLA-B*08:01 (34.4% patient allele frequency vs. 11.44% reference, 3.00-fold; carriers 9 of 16 = 56%; q = 0.019) and HLA-B*40:01 (21.9% vs. 5.28%, 4.14-fold; carriers 7 of 16 = 44%; q = 0.020). HLA-C*03:04 showed suggestive enrichment (21.9% vs. 7.49%, 2.92-fold; carriers 7 of 16 = 44%; q = 0.092). HLA-C*07:01 (2.15-fold; carriers 9 of 16 = 56%) and HLA-A*01:01 (1.90-fold; carriers 8 of 16 = 50%) showed concordant directional enrichment that did not survive multiple-testing correction at the per-allele level (q = 0.102 and q = 0.209, respectively). Per-locus omnibus chi-square statistics indicated significant deviation from reference at HLA-B (χ^2^ = 52.9, p < 0.001) and HLA-C (χ^2^ = 30.6, p < 0.001) but not HLA-A (χ^2^ = 17.3, p = 0.10). Notably, 7 of 16 (44%) GAD65^+^ patients carried at least one copy of HLA-A*01:01, HLA-B*08:01, and HLA-C*07:01, consistent with overrepresentation of the ancestral 8.1 haplotype (A*01:01, B*08:01, DRB1*03:01, DQB1*02:01). The expected carrier frequency in US European populations is approximately 12%, and the observed 3.7-fold enrichment was highly significant (binomial p = 0.0013).

**Table 2.** HLA class I allele frequencies in GAD65^+^ patients compared to reference population.

| HLA allele | Patient n (chromosomes) | Carrier freq. (n/16) | NMDP ref freq. | Fold change | $\chi^2$ | Raw p | BH q |
| --- | --- | --- | --- | --- | --- | --- | --- |
| **HLA-A locus** (omnibus $\chi^2 = 17.3$, $p = 0.10$): | | | | | | | |
| A*01:01 | 10/32 (31.2%) | 8/16 (50.0%) | 16.5% | 1.90 | 5.09 | 0.0317 | 0.209 |
| A*02:01 | 8/32 (25.0%) | 7/16 (43.8%) | 27.6% | 0.91 | 0.10 | 0.8452 | 1.000 |
| A*11:01 | 4/32 (12.5%) | 3/16 (18.8%) | 6.1% | 2.05 | 2.30 | 0.1278 | 0.469 |
| A*68:01 | 3/32 (9.4%) | 3/16 (18.8%) | 3.2% | 2.94 | 3.96 | 0.0811 | 0.446 |
| A*03:01 | 1/32 (3.1%) | 1/16 (6.2%) | 14.0% | 0.22 | 3.14 | 0.1191 | 0.469 |
| A*24:02 | 1/32 (3.1%) | 1/16 (6.2%) | 8.5% | 0.37 | 1.18 | 0.5183 | 0.950 |
| **HLA-B locus** (omnibus $\chi^2 = 52.9$, $p < 0.001$): | | | | | | | |
| **B\*08:01** | **11/32 (34.4%)** | **9/16 (56.2%)** | **11.5%** | **3.00** | **16.61** | **0.0006** | **0.019 \*\*** |
| **B\*40:01** | **7/32 (21.9%)** | **7/16 (43.8%)** | **5.3%** | **4.14** | **17.62** | **0.0012** | **0.020 \*\*** |
| B*07:02 | 3/32 (9.4%) | 3/16 (18.8%) | 13.1% | 0.72 | 0.38 | 0.7922 | 1.000 |
| B*57:01 | 2/32 (6.2%) | 2/16 (12.5%) | 3.7% | 1.71 | 0.62 | 0.3269 | 0.796 |
| B*15:01 | 2/32 (6.2%) | 2/16 (12.5%) | 6.1% | 1.03 | 0.00 | 0.7208 | 1.000 |
| **HLA-C locus** (omnibus $\chi^2 = 30.6$, $p < 0.001$): | | | | | | | |
| C*07:01 | 11/32 (34.4%) | 9/16 (56.2%) | 16.0% | 2.15 | 8.04 | 0.0123 | 0.102 |
| **C\*03:04** | **7/32 (21.9%)** | **7/16 (43.8%)** | **7.5%** | **2.92** | **9.56** | **0.0084** | **0.092 \*** |
| C*07:02 | 3/32 (9.4%) | 3/16 (18.8%) | 14.1% | 0.66 | 0.60 | 0.6131 | 1.000 |
| C*01:02 | 3/32 (9.4%) | 3/16 (18.8%) | 3.4% | 2.75 | 3.46 | 0.0946 | 0.446 |
| C*06:02 | 2/32 (6.2%) | 2/16 (12.5%) | 9.3% | 0.67 | 0.36 | 0.7647 | 1.000 |

**Figure 2.**
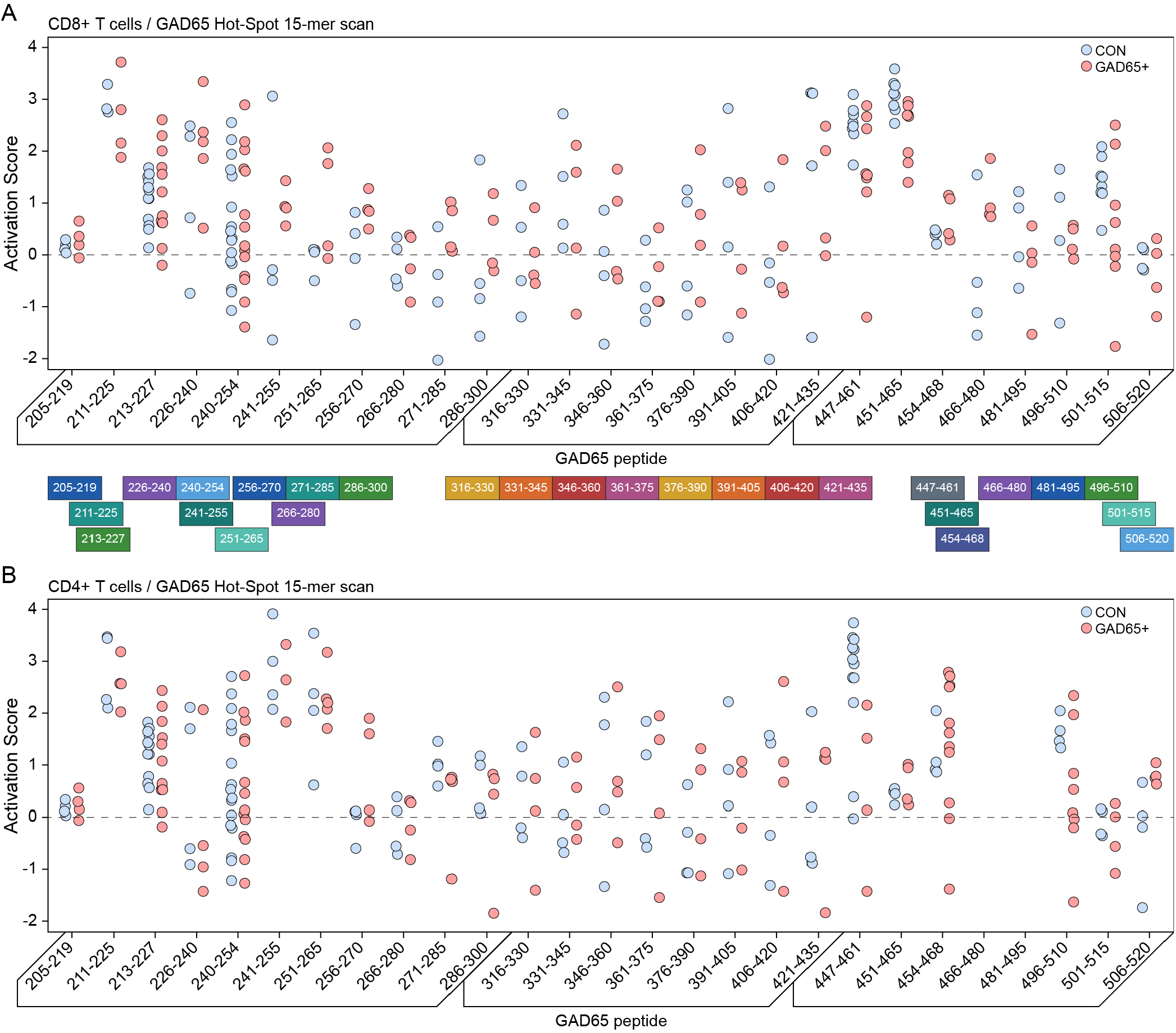
Fine mapping of GAD65 T cell reactivity with individual 15-mer peptides identifies multiple CD8^+^ and CD4^+^ hotspot regions. **A**) CD8^+^ AIM activation scores in response to individual overlapping 15-mer peptides selected from candidate immunogenic regions of GAD65. Each point represents one subject tested against the indicated peptide; blue circles, controls; red circles, GAD65^+^ subjects. Dashed horizontal lines indicate an activation score of 0. Colored boxes below the plot indicate the parent hotspot regions from which the individual 15-mers were selected. Multiple peptides within GAD65_(205–300)_, GAD65_(316–435)_, and GAD65_(447–520)_ elicited CD8^+^ T cell activation in both controls and GAD65^+^ subjects. Within these broader regions, some subdomains, including GAD65_(241–255)_ and GAD65_(251–265)_, showed a response pattern skewed toward GAD65^+^ subjects, whereas others, such as GAD65_(451–465)_, induced stronger responses in controls. **B**) CD4^+^ AIM activation scores to the same panel of individual GAD65 15-mer peptides. As in panel A, each point represents one subject, and the dashed line indicates an activation score of 0. CD4^+^ T cell responses identified partially overlapping but distinct hotspot regions within GAD65, consistent with differences in antigen processing and epitope recognition between CD4^+^ and CD8^+^ T cell compartments. Peptide labels indicate GAD65 residue ranges.

**Figure 3.**
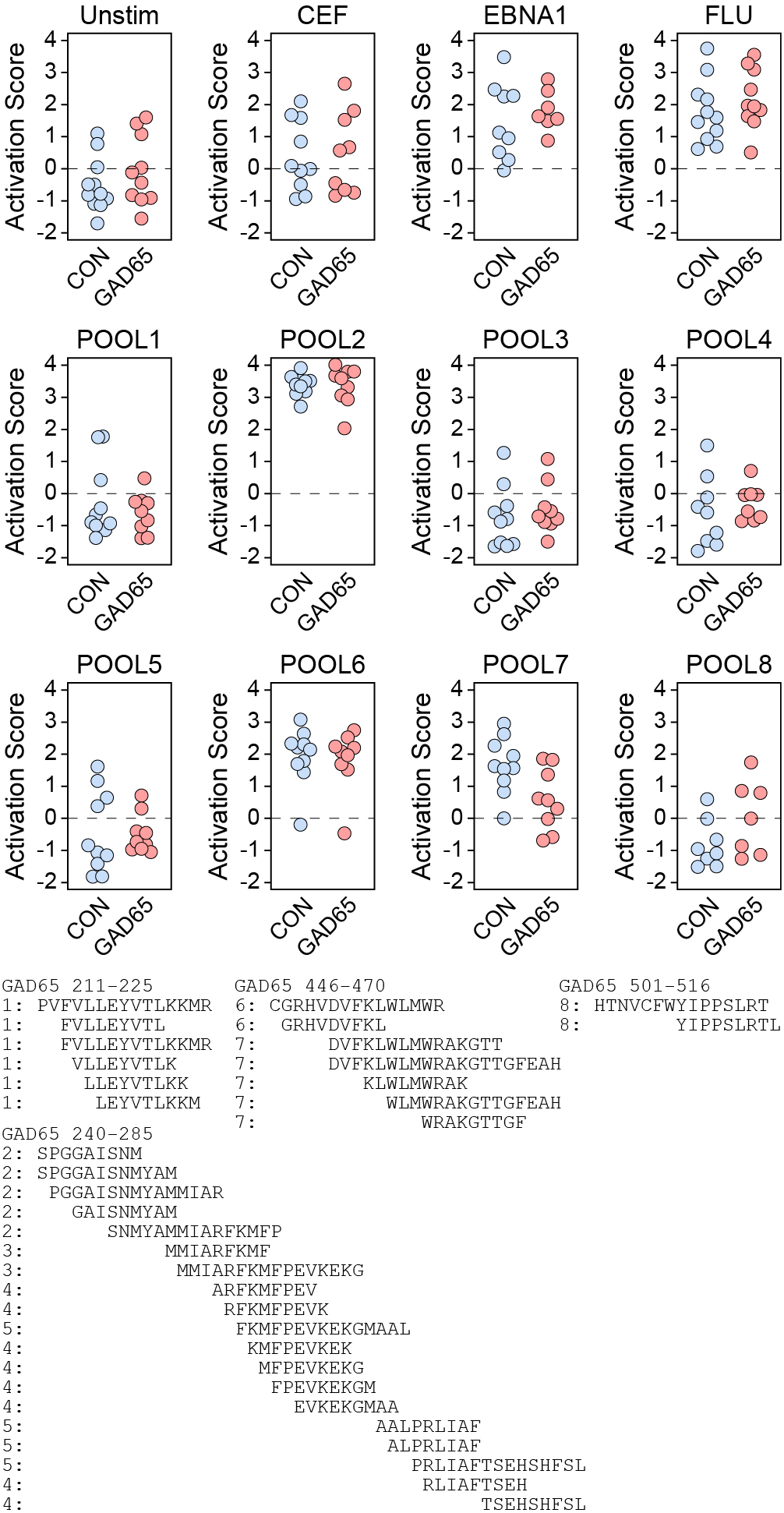
Focused peptide-pool mapping of GAD65 identifies immunogenic subdomains and a region enriched for responses in GAD65^+^ subjects. AIM responses in CD8^+^ T cells to selected GAD65 peptide sub-pools and comparator antigens. Each panel shows activation scores for individual subjects in control (blue) and GAD65^+^ (red) groups; dashed horizontal lines indicate an activation score of 0. Pool composition is shown below the plots, including peptide residue ranges and constituent 9-20-mer sequences. Refinement of the GAD65_(211–225)_, GAD65_(240–285)_, GAD65_(446–470)_, and GAD65_(501–516)_ regions identified two subdomains, POOL2 and POOL6, that induced strong activation in both healthy controls and GAD65^+^ subjects, consistent with broadly immunogenic regions. Other pools showed little or inconsistent activation despite overlapping sequence content, indicating that peptide length, overlap, and local sequence context influenced CD8^+^ T cell recognition.

### Identification of HLA-restricted immunodominant CD8^+^ epitopes in GAD65

A class I peptide “hotspot” map was generated for GAD65 using NetMHCpan to predict binding of 9-11-mers to the HLA allotypes present in our GAD65^+^ patient cohort (Table 2). The per-residue score representing the relative local density of HLA class I binding peptides revealed several peaks (Figure 4A). We combined this information with the pool-based findings and tested patient and control AIM responses to 9-mer peptides in 6 domains across GAD65 (Figure 4B). Using a threshold of 1.5 times the median absolute deviation (MAD) from the median of the control responses for each peptide, the proportion of subjects with an activation score above the threshold for any peptide within the domain was calculated (Figure 4B). Peptides within GAD65_(137-149)_, GAD65_(205-224)_, and GAD65_(240-251)_ induced the largest proportion of above threshold responses in GAD65^+^ subjects. The AIM activation scores induced in CD8^+^ T cells by peptides within these domains (a-k in Figure 4) were heatmapped for the subjects in our cohort with known HLA haplotypes (GAD01, GAD02, GAD04, GAD05, GAD13, GAD15, GAD16, CON03, CON04, CON06) and ranked into responders and non-responders based on the 1.5 x MAD threshold in controls (Figure 4C). Peptide reactivity was heterogeneous across subjects, with a subset of GAD65^+^ patients showing coordinated responses across multiple adjacent peptides, rather than isolated single-peptide activity. GAD01, GAD02, and GAD16 showed the strongest and broadest positive responses across the displayed peptide set, with GAD13 showing a more intermediate pattern. GAD04 and GAD05 were low or negative across the same peptides and GAD15 exhibited no response to any of the peptides. Such clustered, multi-peptide reactivity in a subset of patients suggests coordinated recognition of adjacent epitopes within this region of GAD65. However, peptide responsiveness was not unique to the GAD65^+^ group, as two control subjects also showed multi-peptide positivity, albeit at lower levels that did not cross the threshold as responders. Finally, we mapped the responder status to HLA allotype but observed no specific enrichment in responders, though HLA-A*01:01, HLA-B*08:01, and HLA-C*07:01 were only observed in GAD65^+^ subjects and not in controls (Figure 4C).

**Figure 4.**
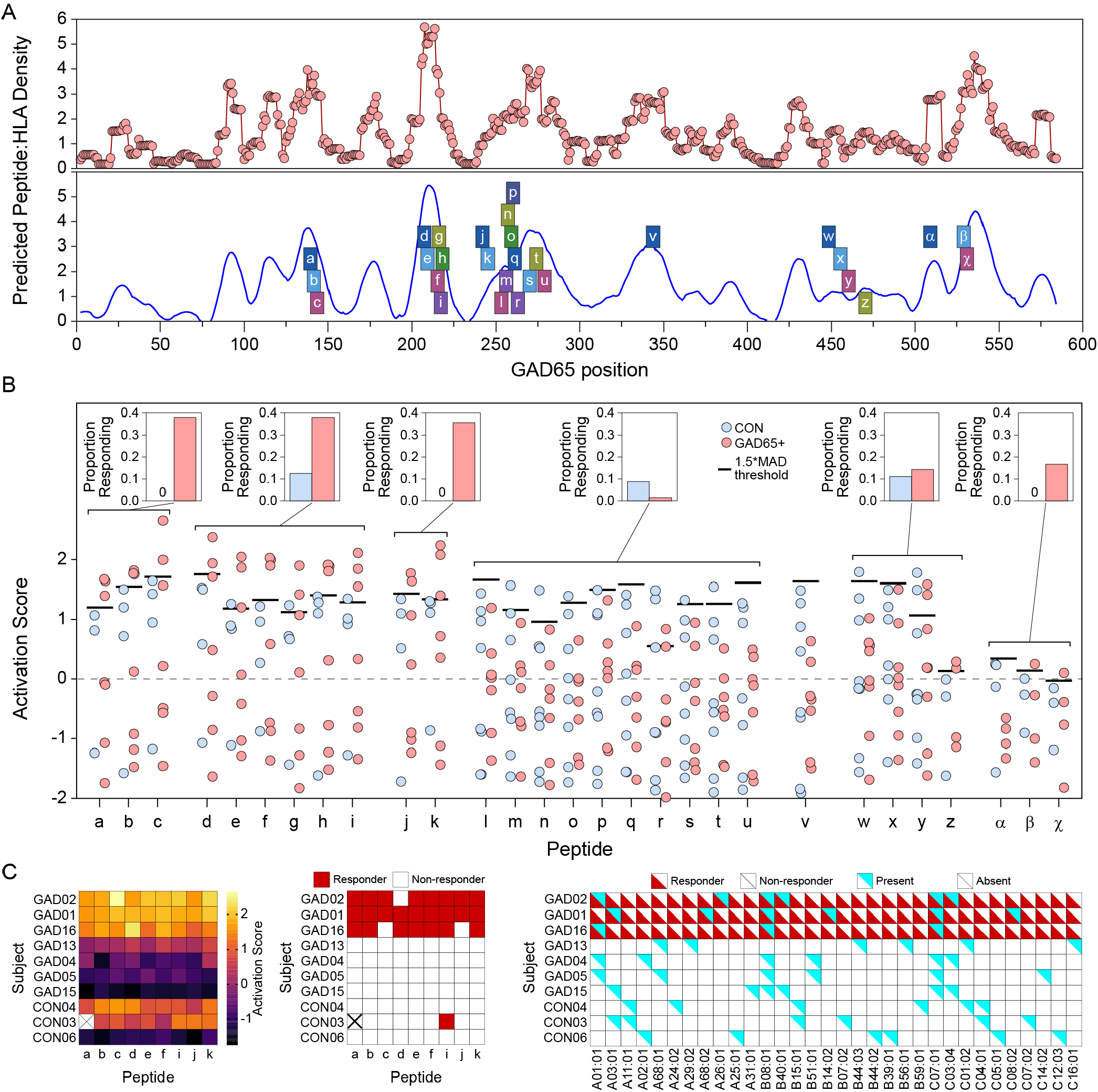
Identification of immunodominant GAD65 class I peptide regions and subject-level CD8 T-cell responses. **A**) A class I peptide “hotspot” map of human GAD65 generated using NetMHCpan predictions for 9-11-mer peptide binding to the HLA class I allotypes represented in the GAD65^+^ cohort. The upper trace shows the per-residue density of predicted peptide:HLA binding across the GAD65 sequence, and the lower trace highlights the principal hotspot peaks selected for downstream testing. Colored boxes indicate the locations of the individual 9-mer peptides and peptide:HLA complexes evaluated experimentally. **B**) CD8^+^ AIM responses to individual 9-mer peptides spanning six candidate domains of GAD65 in GAD65^+^ subjects and controls. Each point represents the activation score for a single subject-peptide pair; black horizontal bars indicate the responder threshold for each peptide, defined as the control median ^+^ 1.5 × median absolute deviation (MAD). Insets show the proportion of subjects with responses above threshold within each domain. Peptides within GAD65_(137–149)_, GAD65_(205–224)_, and GAD65_(240–251)_ produced the largest proportion of above-threshold responses in GAD65^+^ subjects. Peptide labels correspond to: a : YPNELLQEY, b : NELLQEYNW, c : LLQEYNWEL, d : FTYEIAPVF, e : YEIAPVFVL, f : FVLLEYVTL, g : VLLEYVTLK, h : LLEYVTLKK, i : LEYVTLKKM, j : SPGGAISNM, k : GAISNMYAM, l : MMIARFKMF, m : ARFKMFPEV, n : RFKMFPEVK, o : KMFPEVKEK, p : MFPEVKEKG, q : FPEVKEKGM, r : EVKEKGMAA, s : ALPRLIAFT, t : RLIAFTSEH, u : TSEHSHFSL, v : GAFDPLLAV, w : GRHVDVFKL, x : KLWLMWRAK, y : WRAKGTTGF, z : AHVDKCLEL, α : YIPPSLRTL, β : KVAPVIKAR, χ : APVIKARMM. **C**) Subject-level summary of responses to the dominant peptide cluster (peptides a–k) in individuals with known HLA haplotypes. Left, heatmap of AIM activation scores. Middle, binary responder matrix based on the 1.5 × MAD threshold derived from controls. Right, subject HLA class I genotype matrix. Peptide reactivity was heterogeneous across subjects, with GAD01, GAD02, and GAD16 showing the strongest and broadest multi-peptide responses, GAD13 showing an intermediate pattern, and GAD04, GAD05, and GAD15 showing little or no response. Control subjects showed occasional low-level multi-peptide activity, but these responses generally did not exceed the responder threshold. Mapping responder status to HLA allotypes did not reveal enrichment of a single restricting allele, although HLA-A*01:01, HLA-B*08:01, and HLA-C*07:01 were observed only in GAD65^+^ subjects in this cohort.

To determine whether the candidate peptide regions were recognized as HLA class I-restricted epitopes, we generated peptide:HLA tetramers spanning predicted GAD65 class I hotspots and tested tetramer binding on CD8^+^ T cells from GAD65^+^ subjects and controls. To improve specificity for rare antigen-specific events, each peptide-HLA complex was assembled into two independently tetramerized reagents carrying different fluorophores (APC and BV421), and tetramer-positive events were scored only when both fluorophores were bound. Candidate 9-mer peptides were selected from regions of high predicted peptide:HLA binding density across GAD65 and paired with HLA-A, HLA-B, or HLA-C allotypes represented in the cohort (Figure 5A). Tetramer binding was quantified as a normalized tetramer binding score. Across the full tetramer panel, binding patterns were heterogeneous and many peptide:HLA complexes showed similar signal in GAD65^+^ subjects and controls, indicating that predicted HLA binding alone was not sufficient to define disease-associated recognition. Nevertheless, several tetramers showed a pattern skewed toward increased binding in GAD65^+^ subjects. The clearest examples were observed for A*11:01-restricted VLLEYVTLK_(213-221)_, KMFPEVKEK_(257-265)_, and KVAPVIKAR_(529-537)_, as well as B*08:01-restricted EVKEKGMAA_(261-269)_ and APVIKARMM_(530-538)_. Additional tetramers in the GAD65_(205-269)_ region also showed higher binding in subsets of GAD65^+^ subjects, whereas other complexes, including tetramers derived from the GAD65_(446-462)_ and GAD65_(529-537)_ regions in alternative HLA contexts, were comparable between groups or favored controls. We next examined antigen-specific AIM responses within the subset of subjects in our cohort specifically carrying the B*08:01 (Figure 5B) or A*11:01 backgrounds (Figure 5C). Among HLA-B*08:01 subjects, CD8^+^ T cell activation to whole GAD65 was variable but was accompanied by responses to the EVKEKGMAA_(261–269)_ and APVIKARMM_(530–538)_ peptides, whereas controls were all essentially non-responsive. Similarly, among HLA-A*11:01 subjects, responses to whole GAD65 were paralleled by peptide-specific responses to VLLEYVTLK_(213–221)_, KMFPEVKEK_(257–265)_, and KVAPVIKAR_(529–537)_, again with greater activation in GAD65^+^ subjects relative to controls. Within the HLA-B*08:01 restriction, three overlapping 9-mers in the GAD65_(259-276)_ region showed patient-skewed binding (Figure 5A), suggesting a multi-peptide immunodominant region rather than a single dominant epitope. HLA-B*40:01_(206-214)_ tetramer binding was also higher in the four B*40:01-carrying GAD65+ patients than in the single B*40:01-carrying control, paralleling the enrichment of B*40:01 in the cohort haplotyping analysis (Table 2). These data nominate a small set of candidate class I GAD65 epitopes, particularly within the GAD65_(213–221)_, GAD65_(257–276)_, and GAD65_(529–538)_ regions, that are consistently associated with disease-skewed CD8^+^ T cell recognition in the context of selected HLA alleles.

**Figure 5.**
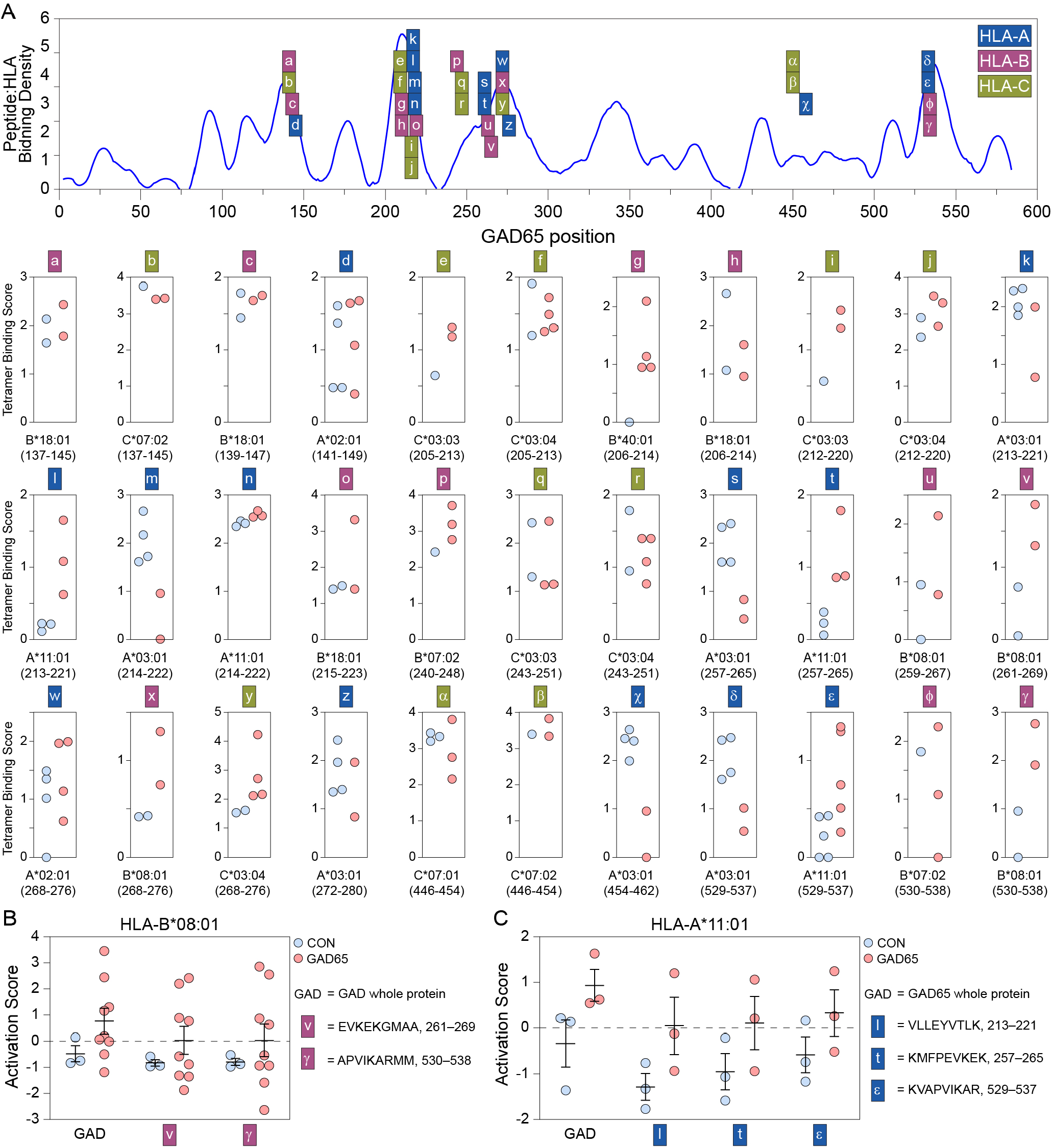
Peptide:HLA tetramer binding on CD8^+^ T cells identifies candidate class I GAD65 epitopes with disease-specificity in selected HLA contexts. **A**) Predicted peptide:HLA class I binding density across the GAD65 sequence (from Figure 4), with positions of the peptide:HLA tetramers that were tested. Label color denotes restricting locus: HLA-A (blue), HLA-B (magenta), and HLA-C (green). Below, each small panel shows the tetramer binding score for one peptide:HLA complex in controls (blue) and GAD65^+^ subjects (red). Peptides are labeled by residue range beneath each panel. Tetramer binding was heterogeneous across the panel, with many candidate complexes showing comparable binding in both groups, but several peptide:HLA combinations showed greater binding in subsets of GAD65^+^ subjects, particularly within the 213-221, 257-269, and 529-538 regions of GAD65. **B**) CD8^+^ AIM activation scores in the subset of subjects carrying HLA-B*08:01, comparing responses to whole GAD65 protein with responses to the EVKEKGMAA_(261–269)_ and APVIKARMM_(530–538)_. **C**) CD8^+^ AIM activation scores in the subset of subjects carrying HLA-A*11:01, comparing responses to whole GAD65 protein with responses to the VLLEYVTLK_(213–221)_, KMFPEVKEK_(257–265)_, and KVAPVIKAR_(529–537)_. Each point in the graphs represents one subject; bars indicate mean ± 95%; dashed horizontal lines indicate an activation score of 0.

### CD8^+^ T cells from GAD65^+^ subjects mediate HLA-restricted killing of target cells expressing GAD65

To determine if the GAD65-reactive CD8^+^ T cell responses identified by AIM and tetramer analyses were associated with direct cytotoxic effector function, we used an HLA reconstitution target-cell system based on MHC class I-deficient HEK-293T cells. Target cells were engineered to express GAD65 together with Cre recombinase and then transduced with Cre-dependent HLA-B08:01-eGFP or control constructs. When cocultured with CD8^+^ T cells from HLA-B*08:01^+^ GAD65^+^ subjects, GAD65-expressing HLA-B*08:01-expressing targets were selectively depleted, whereas this effect was not observed in GAD65-negative targets or HLA-deficient targets or in cocultures with CD8^+^ T cells from HLA-B*08:01^+^ healthy controls (Figure 6A). We further confirmed that T cells from the individuals tested in the HEK killing assay were activated by the candidate HLA-B*08:01-restricted GAD65 peptides EVKEKGMAA_(261–269)_ and APVIKARMM_(530–538)_ (Figure 6B), suggesting that in at least a subset of the GAD65^+^ subjects these were pathogenically relevant immunodominant domains. Live-cell imaging of GAD65-expressing HLA-B08:01-reconstituted HEK targets confirmed that target-cell loss developed progressively after addition of patient CD8^+^ T cells, whereas no-T-cell controls and healthy control cocultures remained stable (Figure 6C). The magnitude of target-cell loss varied among patient donors, with only patient GAD02 exhibiting robust target cell killing, indicating heterogeneity in cytotoxic potential despite shared serologic disease status and HLA haplotype. To determine whether this phenomenon extended beyond B08:01, we tested reconstitution with additional HLA class I alleles represented in the cohort. T cell-dependent loss of GAD65-expressing target cells was observed in a subset of matched HLA contexts across HLA-A, -B, and -C molecules (Figure 6D). These findings indicate that GAD65^+^ subjects harbor functionally cytotoxic CD8^+^ T cells capable of HLA-restricted recognition and elimination of GAD65-expressing target cells, particularly in the context of HLA-B*08:01 and candidate epitopes in the GAD65_(261-269)_ and GAD65_(530-538)_ regions.

**Figure 6.**
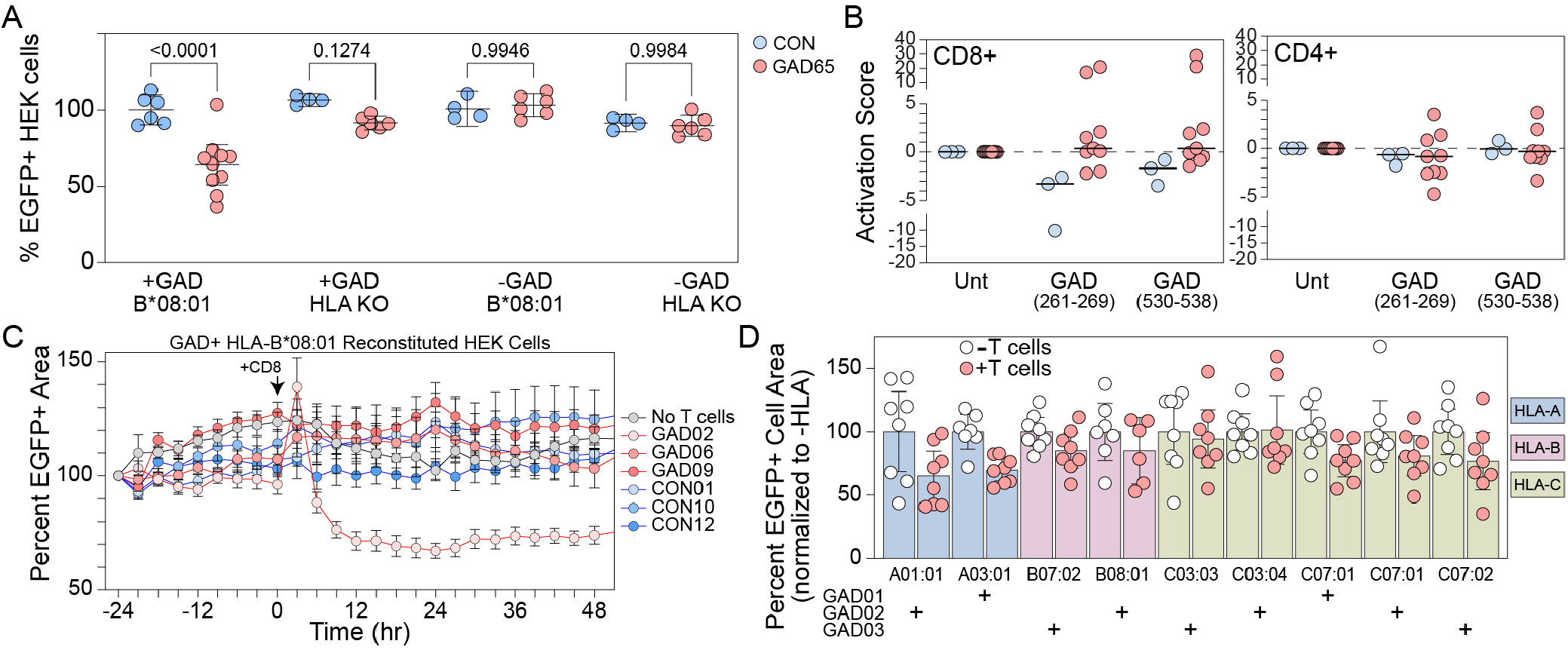
CD8^+^ T cells from GAD65^+^ subjects kill HLA-restricted target cells expressing GAD65. **A**) HLA class I-deficient HEK-293T cells were engineered to express GAD65 and Cre or Cre only and were then transduced with AAV vectors encoding Cre-dependent HLA-B*08:01 and EGFP or EGFP only. After co-culture with CD8^+^ T cells from HLA-B*08:01-positive GAD65^+^ patients (red circles) or healthy controls (blue circles) the percentage of EGFP^+^ target cells was quantified by flow cytometry and normalized to the no T cell condition. Each circle represents one donor. Significance was assessed by two-way ANOVA with Sidak’s pairwise comparison. Bars represent mean ± 95%CI. **B**) Subjects from (A) were tested in the AIM assay to measure activation responses to GAD65 peptides EVKEKGMAA_(261–269)_ and APVIKARMM_(530–538)_. Responses are shown relative to unstimulated controls. Each circle represents an individual healthy control (blue circle) or GAD65^+^ subject (red circle). Black bars represent the median activation score. **C**) Representative live-cell imaging analysis of HEK cells reconstituted with HLA-B*08:01 as in (A) and co-cultured with CD8^+^ T cells from 3 GAD65^+^ patients or 3 healthy control subjects. CD8^+^ T cells were added at t=0. Values are normalized to the GFP^+^ area at t=-24. Data are shown as mean ± 95%CI from at least 6 replicates per condition. **D**) Live-cell imaging of HEK cells reconstituted with HLA-A*01:01, HLA-A*03:01, HLA-B*07:02, HLA-B*08:01, HLA-C*03:03, HLA-C*03:04, HLA-C*07:01, or HLA-C*07:02 and co-cultured with CD8^+^ T cells from the appropriate HLA matched GAD65^+^ subject (GAD01, GAD02, GAD03). Data are mean ± SEM from a minimum of 6 replicates and are normalized to the corresponding HLA-negative condition. Each allotype is shown as replicate wells for the no T cell (open circle) and ^+^CD8^+^ T cell (red circle) condition. Bars are color-coded by HLA.

## Discussion

The immunopathogenesis of GAD65 antibody-associated disorders is incompletely defined, in particular with respect to whether GAD65 antibodies themselves participate in tissue injury or whether they are biomarkers of a broader, T cell-driven autoimmune process directed at an intracellular antigen ^16, 24, 29, 38^. Here we provide evidence for the latter view in a subset of patients with neurological manifestations of GAD65 autoimmunity.First, full-length GAD65, and to a lesser extent GAD67, elicits CD69 upregulation on circulating CD8^+^ and CD4^+^ T cells from GAD65 antibody-positive patients, with CD8^+^ activation in response to GAD65 reaching statistical significance relative to controls. Second, systematic mapping of GAD65 with overlapping 15-mer pools and individual peptides identifies discrete CD8^+^ immunogenic regions that converge with NetMHCpan-predicted class I hotspots and with HLA class I alleles enriched in the patient cohort, including HLA-A*01:01, HLA-B*08:01, and HLA-C*03:04. Third, peptide:HLA tetramers built from candidate hotspot 9-mers detect tetramer-positive CD8^+^ T cells in selected HLA contexts at higher frequency in GAD65^+^ subjects than in controls, and CD8^+^ T cells from a subset of patients mediate antigen- and HLA-restricted killing of GAD65-expressing target cells reconstituted with matched HLA class I alleles.

These observations extend prior reports of CD8^+^ T cell infiltrates in surgical and autopsy material from GAD65-associated temporal lobe epilepsy and limbic encephalitis ^28, 29, 39^, in which perforin-expressing CD8^+^ T cells appose neurons and correlate with neurodegeneration. They also parallel earlier work in T1D demonstrating HLA-A*02:01-restricted GAD65_(114-123)_-specific CD8^+^ T cell clones capable of killing human islet cells ^31^ and HLA class I tetramer-positive GAD65-specific CD8^+^ T cells in T1D peripheral blood ^30^. Together with that literature, the present data support a model in which class I-restricted, GAD65-specific CD8^+^ T cells are part of the immune repertoire engaged in GAD65 autoimmunity and not unique to the pancreatic compartment, although a direct comparison of CNS-targeted and pancreatic-targeted CD8^+^ responses against shared GAD65 epitopes will require dedicated cohorts that include both clinical phenotypes.

Our analysis points to a distinct HLA architecture. Although GAD65 autoimmunity has historically been linked most strongly to class II haplotypes ^14, 25, 40^, consistent with a role for CD4^+^ T cell help and antibody class switching, our data point to a parallel class I axis. Three lines of evidence converge here: cohort-wide enrichment of HLA-B*08:01 and HLA-B*40:01 in non-Hispanic White GAD65^+^ subjects relative to the USA NMDP European Caucasian reference panel (Table 2), with HLA-C*03:04 showing suggestive enrichment in the same direction; a NetMHCpan-derived class I peptide-binding hotspot map that identifies regions of GAD65 likely to be presented across these allotypes; and empirical demonstration that 9-mers drawn from those hotspots elicit AIM responses and tetramer binding skewed toward GAD65^+^ subjects in HLA-A*11:01- and HLA-B*08:01-restricted contexts. The clearest peptide-level signals lie in three regions: GAD65_(213-221)_, GAD65_(257-276)_, and GAD65_(529-537)_. These regions are non-overlapping with the canonical N-terminal humoral epitopes that distinguish GAD65 antibodies in neurological versus T1D contexts ^15, 16^, suggesting that the dominant CD4-supported antibody specificities and the dominant CD8-presented epitopes need not coincide.

The co-enrichment of HLA-A*01:01, HLA-B*08:01, and HLA-C*07:01 in our cohort is consistent with overrepresentation of the ancestral 8.1 haplotype (HLA-A*01:01, HLA-B*08:01, HLA-DRB1*03:01, HLA-DQB1*02:01), one of the most evolutionarily conserved haplotypes in European-descent populations and a well-established risk haplotype for multiple autoimmune diseases, including type 1 diabetes, systemic lupus erythematosus, myasthenia gravis, autoimmune hepatitis, and several thyroid autoimmune conditions ^41^. In our cohort, 44% of non-Hispanic White GAD65^+^ patients (7 of 16) carried at least one copy of all three class I 8.1 alleles, compared to an expected carrier frequency of approximately 12% in US European populations (binomial p = 0.0013). All seven of these patients also carried HLA-DRB1*03:01 and HLA-DQB1*02:01, confirming inheritance of the full 8.1 haplotype rather than coincidental co-carriage of the constituent class I alleles. Two patients (AD016, AD023) carried the full haplotype on both chromosomes, a rate of homozygosity substantially higher than the approximately 0.4% expected in US European populations. Of the two Hispanic/Latino patients excluded from the HLA enrichment analysis, one (AD009) also carried the full 8.1 haplotype, suggesting that this association is unlikely to be confined to non-Hispanic White GAD65^+^ patients. Recognition that GAD65 autoimmunity associates with the 8.1 haplotype links this disease to the broader landscape of autoimmunity driven by impaired tolerance in carriers of this ancestral haplotype, and suggests that the CD8^+^ T cell responses we describe may operate within a permissive immunogenetic background rather than reflecting a GAD65-specific HLA association. Separately, the B*40:01 and C*03:04 co-enrichment we observe is consistent with a second class I risk haplotype distinct from 8.1, as B*40:01 and C*03:04 are in linkage disequilibrium in European populations. Larger cohorts with full haplotype-level phasing will be needed to test whether these two HLA architectures represent independent risk pathways or convergent presentation of overlapping GAD65 epitope domains.

Cohort-wide differences between GAD65^+^ subjects and controls in the magnitude of peptide-specific CD69 upregulation were modest and heterogeneous. Several control subjects also showed peptide-level activation, albeit usually below the control-derived 1.5xMAD threshold. This is consistent with prior reports that low-frequency GAD65-reactive CD8^+^ T cells circulate as a “background” population in healthy individuals ^42^. Rather than a single dominant public epitope, our subject-by-peptide matrices show clustered, multi-peptide reactivity in a subset of GAD65^+^ subjects, with other GAD65^+^ subjects showing no detectable response. This subject-level heterogeneity is a major caveat against treating mere detection of GAD65-specific CD8^+^ T cells as evidence of pathogenicity, and echoes observations in organ-specific autoimmunity ^43^ in which qualitative features of autoreactive T cells such as differentiation state, cytotoxic potential, tissue access, and metabolic programming are likely to determine whether a circulating clone contributes to target organ injury.

The HLA-restricted HEK target-killing assay (Figure 6) links antigen recognition to cytotoxic effector function but should be read with appropriate caution. Target-cell loss in the HLA-B*08:01 reconstitution experiment required both GAD65 expression and matched HLA-B*08:01; however, the magnitude of killing varied substantially across patient donors sharing serologic disease status and HLA-B*08:01, with robust killing observed in only one donor (GAD02) in the dedicated B*08:01 experiment. The expanded HLA panel (Figure 6D) likewise showed donor- and allele-specific patterns, with each HLA allele tested in a single matched patient donor. Notably, GAD02 itself showed differential responses across its own HLA alleles, with stronger killing of HLA-A*01:01-reconstituted than HLA-B*08:01-reconstituted targets in the cross-allele panel, consistent with a polyclonal autoreactive repertoire in which different class I alleles dominate antigen presentation in different patients. These data indicate that GAD65-specific, HLA-restricted cytotoxic activity exists in at least a subset of GAD65^+^ subjects and across multiple HLA class I contexts, but its magnitude is heterogeneous, donor-specific, and not predicted by HLA haplotype alone. Whether this heterogeneity reflects differences in clonal frequency, T cell differentiation state, exhaustion, ongoing immunotherapy, or variability in the engineered target system will require larger HLA-stratified cohorts and direct phenotyping of the responding clones.

Several limitations should be made explicit. First, the cohort is modest in size and clinically heterogeneous, spanning SPS, cerebellar ataxia, and epilepsy; we did not stratify analyses by clinical phenotype, and additional subjects in each phenotype will be needed to determine whether the immunological signatures defined here vary by syndrome. Second, the GAD65^+^ patient cohort skews older than the healthy control cohort, and age-related shifts in T cell repertoire diversity, naive/memory composition, and inflammatory tone could contribute to the patient-vs-control differences observed in the AIM assays; sensitivity analyses adjusting for age, and recruitment of additional age-matched controls, will be important to confirm that the differences described here reflect disease rather than demographics. Third, immunotherapy histories at the time of sampling were variable, with most patients receiving some form of immunomodulation within the 6 months preceding PBMC collection (Table 1); within the limits of a small untreated subset, peripheral CD8 activation scores were comparable between untreated patients and those receiving immunotherapy, indicating that treatment is unlikely to be the primary driver of the patient-vs-control differences shown in Figure 1, although larger treatment-stratified cohorts will be needed to formally exclude treatment-driven effects on more sensitive readouts. Fourth, our immunological assays are anchored on an early activation marker (CD69), and we did not systematically profile exhaustion, tissue homing, or cytotoxic effector molecules that would distinguish pathogenic effectors from bystanders. Direct phenotyping of tetramer-enriched CD8^+^ T cells, including granule content (granzyme B, perforin), differentiation state (CCR7, CD45RA), exhaustion (PD-1, TIGIT), and degranulation capacity (CD107a), is a priority for our ongoing work. Fifth, the HEK-293T reconstitution assay isolates the antigen-HLA-TCR interaction from the broader CNS milieu, including glial costimulation, the chronic inflammatory environment, and barrier biology. Sixth, the per-allele n in Figure 6D is small, and the most striking single effect is driven by a single patient. Replication across additional HLA-matched donors will be required before any specific class I allele can be designated as a clinically actionable target. Seventh, the cross-sectional design precludes inferences about temporal evolution of GAD65-specific T cell repertoires, including their response to specific immunotherapies. Eighth, HLA enrichment analyses were restricted to the 16 non-Hispanic White GAD65^+^ patients to ensure ancestral compatibility with the European Caucasian reference panel; whether the immunogenetic associations described here generalize to GAD65^+^ patients of non-European ancestry will require larger, ancestrally diverse cohorts and ancestry-matched reference populations. Finally, we have not directly demonstrated cytotoxicity against human neurons; that experiment remains an important next step.

In summary, this study integrates HLA class I typing, NetMHCpan-based hotspot prediction, peptide:HLA monomer affinity testing, AIM-based functional mapping, peptide:HLA tetramer staining, and HLA-reconstituted target-cell killing in a single cohort of patients with GAD65 antibody-associated neurological disorders. We identify GAD65_(213-221)_, GAD65_(257-276)_, and GAD65_(529-537)_ as candidate immunodominant CD8^+^ epitopes restricted by HLA-A*11:01 and HLA-B*08:01, demonstrate enrichment of tetramer-positive CD8^+^ T cells against these epitopes in GAD65^+^ subjects, and provide evidence that at least some GAD65^+^ patients harbor CD8^+^ T cells capable of HLA-restricted cytotoxicity against GAD65-expressing target cells. We further show enrichment of the autoimmunity-associated 8.1 ancestral haplotype in 44% of non-Hispanic White GAD65^+^ patients, linking GAD65 autoimmunity to a class I-restricted immunogenetic background shared with multiple other autoimmune diseases. The principal advance of this work is a panel of validated peptide:HLA tetramers built around GAD65 9-mers presented by HLA alleles enriched in patient cohorts. These reagents enable experiments not previously feasible in this disease: longitudinal tracking of antigen-specific clones in individual patients, single-cell TCR sequencing of tetramer-enriched populations, direct functional and phenotypic comparison of CD8^+^ T cells from patients in different clinical states or on different therapies, and parallel assessment in CSF where the relevant pathogenic populations are likely to be enriched. Together, these data argue for a class I-restricted CD8^+^ T cell arm in GAD65 autoimmunity that has been underrepresented relative to the long-standing focus on antibodies and CD4^+^ help, and they provide reagents and an approach for the prospective studies needed to establish whether GAD65-specific CD8^+^ T cells contribute directly to neuronal injury across the GAD65 antibody-associated neurological disease spectrum.

## AUTHOR CONTRIBUTIONS

CLH and BDSC conceptualized the project and manuscript. PS, BDSC, and CLH drafted the manuscript. PS, BDSC, BLO, RBS, and JC contributed to data collection. BDSC, CLH, and PS designed and drafted the figures. CLH supervised the study, provided overall guidance, and edited and approved the final version of the manuscript. All the authors contributed to review of the manuscript.

## CONSENT FOR PUBLICATION

Not applicable.

## ACKNOWLEDGEMENTS

The authors acknowledge support and resources provided by the Mayo Clinic Center for Multiple Sclerosis and Autoimmune Neurology (CMSAN). We acknowledge our clinical colleagues, including Sean Pittock, Anastasia Zekeridou, Andrew McKeon, and Samantha Banks, for their contributions to patient identification and recruitment. We acknowledge the expert technical assistance of Jonghoon Choi and Ramila Barun Shrestha.

