## Supplemental Table 1 for "Characterization of HLA-restricted GAD65-specific CD8^+^ T cell responses in patients with GAD65 antibody-associated neurological disorders"

**Supplemental Table 1. Control demographics.**

| Participant ID | Age | Sex |
|---|---|---|
| CON01 | 40 | Male |
| CON02 | 37 | Female |
| CON03 | 34 | Female |
| CON04 | 42 | Male |
| CON05 | 30 | Male |
| CON06 | 24 | Female |
| CON07 | 23 | Female |
| CON08 | 21 | Male |
| CON09 | 24 | Male |
| CON10 | 21 | Female |
| CON11 | 45 | Female |
| CON12 | 28 | Female |
| CON13 | 26 | Male |
| CON14 | 33 | Female |
| CON15 | 60 | Male |
